# Contribution of CENP-F to FOXM1-mediated discordant centromere and kinetochore transcriptional regulation

**DOI:** 10.1101/2023.12.27.573453

**Authors:** Sakshi Khurana, Daniel R. Foltz

## Abstract

Proper chromosome segregation is required to ensure genomic and chromosomal stability. The centromere is a unique chromatin domain present throughout the cell cycle on each chromosome defined by the CENP-A nucleosome. Centromeres (CEN) are responsible for recruiting the kinetochore (KT) during mitosis, ultimately regulating spindle attachment and mitotic checkpoint function. Upregulation of many genes that encode the CEN/KT proteins is commonly observed in cancer. Here, we show although that FOXM1 occupies the promoters of many CEN/KT genes with MYBL2, occupancy is insufficient alone to drive the FOXM1 correlated transcriptional program. We show that CENP-F, a component of the outer kinetochore, functions with FOXM1 to coregulate G2/M transcription and proper chromosome segregation. Loss of CENP-F results in alteration of chromatin accessibility at G2/M genes, including CENP-A, and leads to reduced FOXM1-MBB complex formation. The FOXM1-CENP-F transcriptional coordination is a cancer-specific function. We observed that a few CEN/KT genes escape FOXM1 regulation such as CENP-C which when upregulated with CENP-A, leads to increased chromosome misegregation and cell death. Together, we show that the FOXM1 and CENP-F coordinately regulate G2/M gene expression, and this coordination is specific to a subset of genes to allow for proliferation and maintenance of chromosome stability for cancer cell survival.

## Introduction

A hallmark of cancer, chromosome instability (CIN) is the alteration of the number or structure of chromosomes[1, 2]. CIN is a potent source of genetic variation which allows tumors to adapt to new environments, increase proliferation and develop therapeutic resistance[1, 3, 4]. Errors in chromosome segregation are an established source of CIN [2, 5]. Chromosome misegregation events can lead to the formation of micronuclei and result in chromosomal shattering (or chromothripsis) and subsequent rearrangements[6]. It is proposed that precancerous lesions can directly arise from these genomic crises[6, 7].

Periodic changes in gene expression are coordinated with cell cycle progression. Cancer cells often use this coordination to alter cell proliferation and enhance diversity [8]. The DREAM complex, consisting of four core components – dimerization partner proteins (<u>D</u>P), retinoblastoma (<u>R</u>b) family proteins, <u>E</u>2F transcription factors and the <u>M</u>uvB complex, is responsible for timely activation and repression of G1/S and G2/M genes [9, 10]. These genes are discriminated by their promoter sites bound by the DREAM complex; G1/S genes contain E2F elements and are repressed by E2F-DP, while G2/M genes contain CHR sites and are repressed by MuvB. Cell-cycle dependent phosphorylation of DREAM components by Cdk-cyclin proteins results in disruption of this repression[8, 11]. One of the key G1/S genes is MYBL2 which binds MuvB to form the MMB complex[12]. This complex recruits FOXM1 in G2/M[13]. MYBL2 is then degraded, leaving FOXM1 bound to CHR sites with MuvB to activate mitotic genes[10, 14].

Among the G2/M genes, centromere, and kinetochore (CEN/KT) genes are critical for chromosome segregation. During mitosis, replicated genomic DNA condenses into mitotic chromosomes with sister chromatids being attached at the centromere, a highly specialized region of chromatin that specifies the site of kinetochore formation on chromosomes[15]. Centromeres are epigenetically marked by centromere protein A (CENP-A), a histone H3 variant[16]. CENP-A recruits inner kinetochore components which then attach to the microtubule-binding outer kinetochore protein structure, responsible for proper attachment of chromatin to the mitotic spindle[17–19].

A highly regulated process, chromosome segregation is dependent on the CEN/KT and improper function of the complex can lead to misegregation errors [5, 20]. For example, overexpression of CENP-A leads to mislocalization of other CEN/KT components and subsequent mitotic defects[21–23]. CEN/KT gene mis-expression is correlated with poor patient outcomes such as survival and response to radio- and chemotherapy[24]. While studies of overexpression of specific CEN/KT components are informative, cancers are characterized by coordinated misexpression of the whole complex. This suggests the existence of a mitotic transcriptional program that is uniquely mis-regulated in cancers and this coordinated misregulation, may play a role in cancer initiation[24, 25].

Highly aneuploid breast tumors enriched in p53 mutations are co-associated with overexpression of the three cell-cycle transcription factors – FOXM1, E2F1 and MYBL2. Triple overexpression of the transcription factors resulted in increased chromosome segregation defects compared to the single expression cells in a xenopus model system [26]. FOXM1, in particular, is proposed to be the master regulator of mitosis[27]. FOXM1 is overexpressed in many cancers and correlated with tumor progression [28, 29]. It is involved in a variety of cancer-related cell processes including DNA damage repair and chemotherapy resistance[30]. Overexpression of FOXM1 in MEFs leads to increased cell proliferation and mitotic index and depletion results in mitotic failure and chromosomal misalignment[31]. Interestingly, the coordinated upregulation pattern of CEN/KT genes is highly correlated FOXM1 expression[25]. In addition, FOXM1 binds the promoters of several CEN/KT and other mitotic genes including PLK1, CCNB1, CDC25B, AURKB, CENPA, and CENPF [31, 32]. Thus, FOXM1 is suggested to drive the temporal cell-cycle dependent transcription of these G2/M genes including the CEN/KT genes[33]. However, the context specific changes of the FOXM1 transcriptional program in cancer cells compared to normal cells is poorly understood.

CENP-F, a target of FOXM1, is a potential cofactor of the transcriptional regulation. Canonically a microtubule-binding member of the outer kinetochore, CENP-F has been assigned several additional functions [34]. It accumulates during the S/G2 phase at the nuclear matrix and envelope, and during late G2/M and localizes to the centromeres where it plays a critical role in chromosome alignment and segregation[31, 35–37]. Relevantly, a cross species computational analysis of master regulators of prostate cancer (PCa) progression, predicted that CENP-F and FOXM1 co-regulate target genes involved in development of PCa[38]. Functional validation of these two regulators reveals that co-expression is an indicator of poor patient prognosis. Furthermore, co-silencing CENP-F and FOXM1 leads to abrogation of tumor growth as well as deregulation of mitotic and cell-cycle related transcriptional targets[38]. In prostate cancer cells, silencing CENP-F alone leads to mislocalization of FOXM1 and reduced occupancy of FOXM1 at promoters of target genes[38]. CENP-F has previously been implicated in transcriptional regulation. It binds and sequesters retinoblastoma protein (Rb) and is proposed prevent its binding with E2F1, allowing transcription of E2F targets during the G1/S transition[39, 40]. CENP-F also interacts with Activating Transcription Factor 4 (ATF4) via leucine zipper domains[41]. Together, these previous studies provide compelling evidence that CENP-F may play an important role in cell-cycle dependent gene transcription.

Here we show that while FOXM1 is correlated with the CEN/KT genes, overexpression of FOXM1 is insufficient to drive overactivation of the G2/M gene pathway in normal human cells. We show that that CENP-F and FOXM1 regulate similar sets of genes involved in G2/M function and progression. The correlated gene expression can be explained to some degree by the observation that loss of CENP-F changes the chromatin landscape at G2/M genes, including CENP-A, and weakens FOXM1-MMB complex formation. CENP-A is transcriptionally downregulated with loss of CENP-F and FOXM1 in cancer cells but not in normal cells, suggesting that the transcriptional coordination of CENP-F and FOXM1 is functionally distinct in the context of cancer. Furthermore, while FOXM1 and MYBL2 co-occupy promoters and transcriptionally activate many CEN/KT genes, not all genes are controlled as part of this pathway. The selective escape of CENP-C from FOXM1 regulation promotes cell survival. Overall, we determine that CENP-F and FOXM1 coordinate to drive transcriptional activation of a specific subset of G2/M genes in association with the MuvB complex to promote cancer cell proliferation and survival.

## Results

### FOXM1 and CENP-F coordinately regulate mitotic pathways

To determine the necessity of FOXM1 for transcriptional regulation of CEN/KT genes in non-transformed cells and elucidate the role that CENP-F plays in this function, we knocked down CENP-F and FOXM1 in near diploid, and stably karyotypic hTERT-RPE1 cells using siRNA for 48 hours. We observed a significant decrease in levels of the respective proteins in the siCENPF, siFOXM1 and siDouble conditions by immunoblotting (Figure 1A). Interestingly, while protein levels of CENP-A are not fully ablated by any of the conditions, they were mildly reduced in siCENP-F and siFOXM1 conditions and noticeably reduced in the siDouble condition (Figure 1A). Immunofluorescence of the cells with FOXM1 and CENP-F antibodies confirms knockdown (Figures 1B and 1C). Loss of CENP-F and FOXM1 caused an increase in micronuclei formation and lagging anaphase bridges, however, double knockdown of both CENP-F and FOXM1 led to greater chromosome instability (Figure 1D-F). Thus, while loss of either CENP-F or FOXM1 results in chromosome instability phenotypes, there is a combinatorial effect seen with the double knockdown.

**Figure 1.**
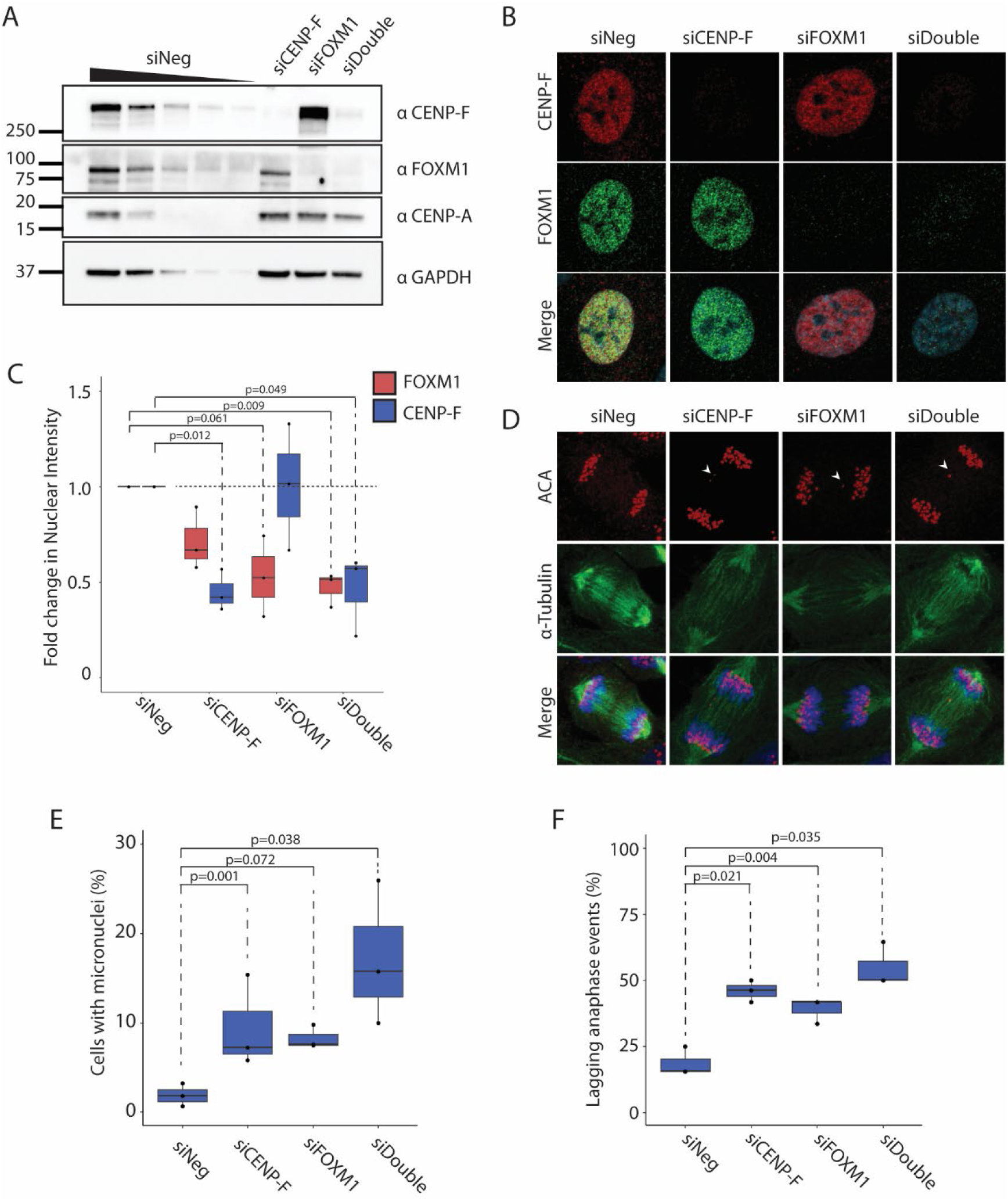
CENP-F and FOXM1 systematically regulate chromatin and centromeric stability. A. Immunoblot analysis showing protein knockdown in response to 20nM siRNA treatment of CENP-F, FOXM1 or double KD for 48 hours. Wedge represents loading titration: 1x > 0.75x > 0.5x > 0.25x > 0.125x. All other lanes are loaded at 1x concentration. N=3. B. Representative images of hTERT-RPE1 cell treated with siCENP-F, siFOXM1 or both. Immunofluorescence was conducted to visualize CENP-F (green) and FOXM1 (red), and DAPI (blue). C. Quantification of CENP-F and FOXM1 protein levels from image set shown in (B). P-values calculated using two-tailed paired student’s t-test. N=3. D. Representative mitotic images of hTERT-RPE1 cell treated with siCENP-F, siFOXM1 or both. Immunofluorescence was conducted to visualize centromeres by ACA (red), alpha-Tubulin (green), and DNA by DAPI (blue). E. Quantification of micronuclei frequency from (C). P-values calculated using two-tailed paired student’s t-test. N=3. F. Quantification of lagging anaphase events from (C). P-values calculated using two-tailed paired student’s t-test. N=3.

To examine the effect of CENP-F and FOXM1 loss, we conducted transcriptome analysis using total RNA-seq. The siRNA treatment resulted in 87.8% loss of CENP-F and 88.7% loss of FOXM1 RNA levels in the individual knockdowns and 82% of each in the double knockdown (Figure 2A). Cells depleted of CENP-F or FOXM1 show a highly correlated pattern of gene expression changes (Figure 2B and 2C). While the degree of knockdown was similar, siRNA mediated knockdown of FOXM1 results in more pronounced downstream effects compared to CENP-F as indicated by the number of downregulated genes passing the significance threshold (Supplementary Figure 1A). Despite the highly correlated changes, the most highly downregulated genes in each knockdown condition show limited overlap (Supplementary Figure 1B). Centromere and kinetochore genes are marginally downregulated in both conditions (Supplementary Figure 1C).

**Figure 2.**
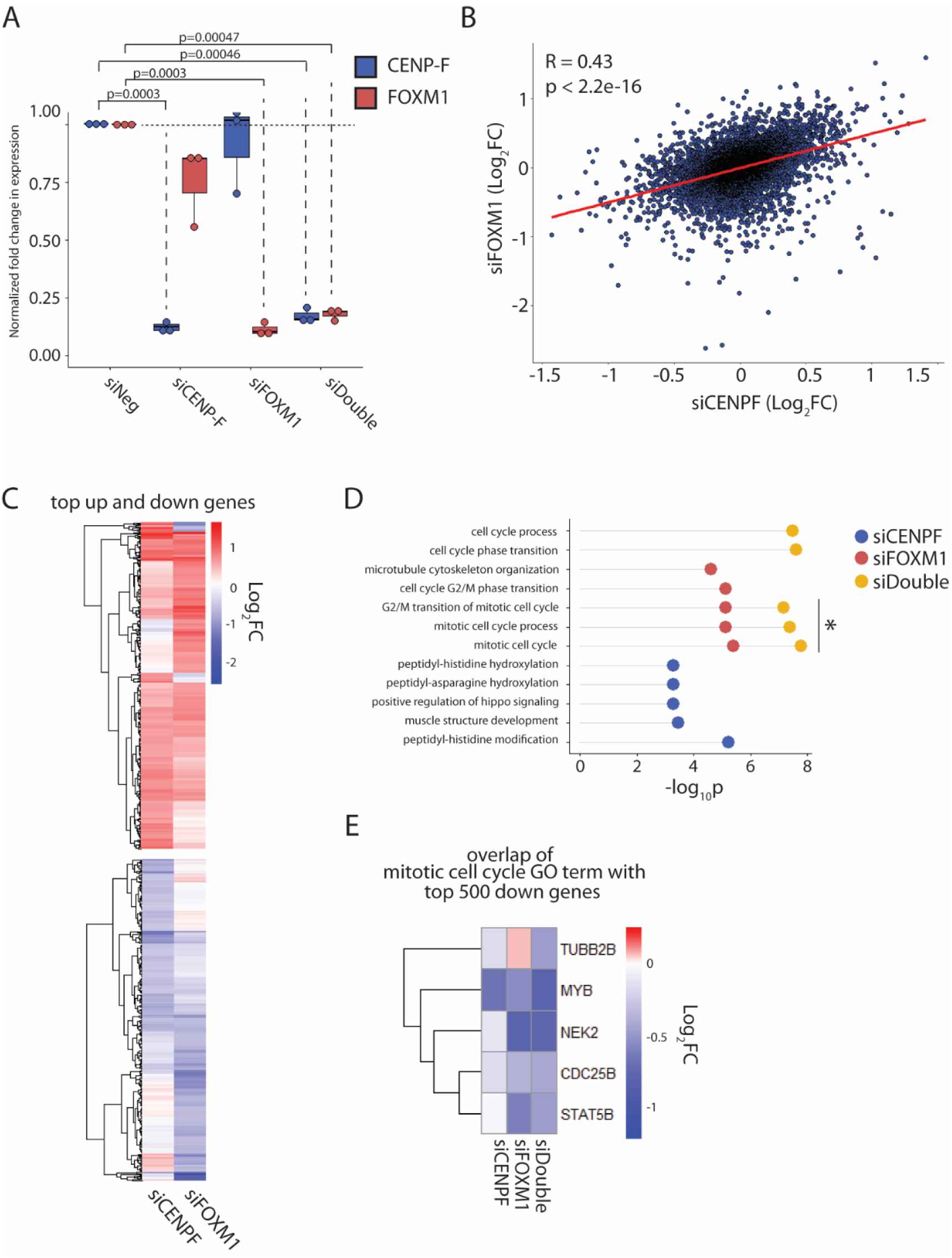
CENP-F and FOXM1 coordinately co-regulate G2/M genes. A. Fold change of CENP-F and FOXM1 transcript levels normalized to GAPDH, in response to 20nM siRNA treatment for 48 hours in hTERT-RPE1 cells. P-values calculated using two-tailed paired student’s t-test. N=3. B. Scatterplot of genes from RNA-seq plotted as log_2_FC in siCENP-F vs siFOXM1 treated cells. Correlation coefficient and p-value was calculated using Pearson method. N=3. C. Heatmap showing log_2_FC of top 500 upregulated and top 500 downregulated genes in siCENP-F and siFOXM1 conditions, clustered by unbiased Euclidean method. D. GO-term analysis showing top 5 enriched GO-terms in top 500 downregulated genes in siCENP-F, siFOXM1 and siDouble cells. GO-terms enriched in multiple conditions are plotted on a single line with different colored circles. E. Heatmap showing overlapping genes between top 500 down genes in the RNA-seq datasets and the mitotic cell cycle gene set (enriched in both siFOXM1 and siDouble). Log_2_FC of overlapping genes are shown in all conditions.

Double knockdown of CENP-F and FOXM1 have a more profound effect than individual knockdown. GO enrichment analysis reveals that within the top five enriched pathways in each condition, the double knockdown effects more mitotic and cell cycle pathways in comparison to either single knockdown as well exhibiting more significant enrichment of the pathways (Figure 2D). Overlap of the top 500 genes in the dataset with genes in one such pathway, mitotic cell cycle (GO:0000278), does not reveal a higher degree of downregulation at the individual gene level in the double knockdown compared to the individual knockdowns (Figure 2E). This suggests that the coordinated function of FOXM1 and CENP-F is systemic regulation of mitotic and cell cycle pathways to support proper chromosome segregation.

### FOXM1 is insufficient to drive the activation of mitotic genes and may require additional factors

Given that FOXM1 has a larger transcriptomic effect compared to CENP-F in our system, we aimed to evaluate the sufficiency of FOXM1 for the upregulation of G2/M genes in normal cells. To examine the effect of FOXM1 overexpression on transcription, we used lentiviral delivery to create doxycycline inducible FLAG-FOXM1c expressing cells and validated the system (Supplemental Figure 2A). Activation of FLAG-FOXM1c was successful in our cell line, however we did not observe upregulation of CENP-A and CENP-F protein levels. We then conducted RNA-seq and found that FOXM1 overexpression did not lead to transcriptomic changes of centromere and kinetochore genes (Supplementary Figure 2B). Instead, upregulated genes include DSC3, SLC43A2 and IGFBP1 among other developmentally important genes (Supplemental Figure 2B). Surprisingly, GSEA reveals that cell cycle related gene sets are downregulated including CDCA3 (Supplemental Figures 2B and C). Indeed, centromere and kinetochore genes are generally downregulated (Supplemental Figure 2D). Further examination of the data using GO term analysis of the top genes reveals that pathways enriched in the significantly upregulated genes (cluster 2) include tissue development and neuroepithelial cell differentiation (Supplementary Figure 2E and F). Surprisingly, the significantly downregulated genes (cluster 1) are enriched in mitotic cell cycle processes and other cell division related pathways (Supplementary Figure 2E and F). Thus, while FOXM1 overexpression may contribute to carcinogenesis through upregulation of developmental and morphogenesis pathways, it is insufficient to drive transcriptional upregulation of CEN/KT genes in normal cells. Instead, FOXM1 overexpression seems to downregulate G2/M genes, possibly a result of a negative feedback loop or its proposed repressive function during interphase[42]. Thus, we hypothesized that additional co-factors are indeed required for the activation of the G2/M pathway.

### Loss of CENP-F leads to changes in chromatin accessibility at G2/M genes bound by FOXM1

FOXM1 occupies promoters to drive transcription of G2/M specific gene. Although CENP-F influences a similar set of genes, how CENP-F exerts this activity is unknown. Immunoprecipitation of FLAG-FOXM1 does not co-purify CENP-F (Supplementary Figure 3A) similar to what has been observed by others [38]. Thus, CENP-F may contribute to G2/M gene regulation independent of binding. We assessed promoter occupancy of FOXM1 and MYBL2 at cell cycle related genes in the presence and absence of CENP-F using ChIP-qPCR. Using AURKB and CCNB1 promoters as known FOXM1 targets, no differences in promoter occupancy were observed in asynchronous cells (Supplementary Figure 3B). Since the association of FOXM1 with MuvB is cell cycle regulated, we enriched the samples for G2/M cells using RO-3306 treatment (Supplementary Figure 3C). While results in RO-3306 treated cells were also statistically insignificant, FOXM1 occupancy trends towards decreasing in response to loss of CENP-F while MYBL2 trends towards increased occupancy at both the AURKB and CCNB1 promoters (Supplemental Figure 3D).

To determine whether CENP-F plays a role at the FOXM1 bound sites in the genome independent of occupancy, we assayed chromatin accessibility changes in response to CENP-F and FOXM1 knockdown by ATAC-seq in asynchronous hTERT-RPE1 cells. Changes in accessibility in both conditions were mapped to FOXM1 ChIP-seq peaks from publicly available data in MCF7 cells and then subjected to k-means clustering (k=3) (Figure 3A). Overall, FOXM1 targets that showed accessibility changes in response to loss of FOXM1 also showed an effect in response to CENP-F loss. Specifically, regions in cluster 1 showed increased accessibility in the absence of either CENP-F or FOXM1. Cluster 3 are regions with little to no change in accessibility in either condition. Most interestingly, cluster 2 included regions differentially affected by CENP-F and FOXM1. FOXM1 loss resulted in decreased accessibility in these regions, consistent with the gene activation function of FOXM1. In contrast, CENP-F loss resulted in increased accessibility in these regions suggesting that CENP-F may normally be bound at those sites.

**Figure 3.**
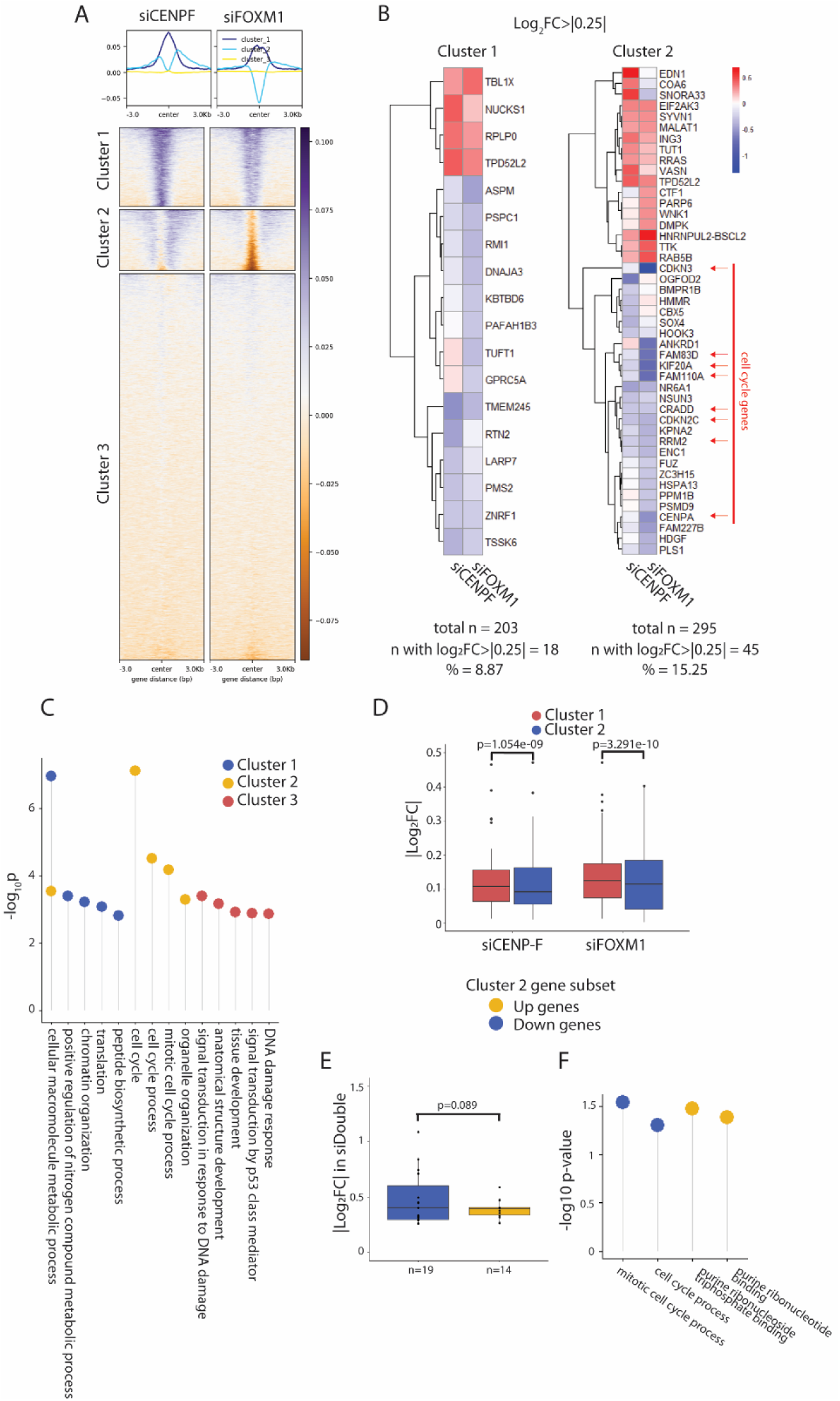
CENP-F influences chromatin landscape of G2/M genes. A. Log_2_FC of ATAC-seq signal in hTERT-RPE1 cells treated with siCENP-F and siFOXM1 for 48 hours, mapped against FOXM1 binding sites in MCF7 cells, with k-mean clustering = 3 (bottom). Enrichment plot shows average accessibility profile in each cluster (top). N=3. B. GO-term enrichment analysis of each cluster. Terms enriched in multiple groups are represented on the same line. Negative log_10_ p value of enrichment is represented on the y-axis. C. Heatmap of the subset of genes in cluster 1 and 2 from ATAC-seq that have log_2_FC>|0.25| in the RNA-seq data set. Cluster 1 has a total of 203 genes with 18 genes with log_2_FC>|0.25| while cluster 2 has a total of 295 genes with 45 genes with log_2_FC>|0.25|. Cell cycle associated genes are highlighted with red arrows. D. Absolute values of log_2_ fold transcriptional changes of downregulated genes in Cluster 1 and Cluster 2 from ATAC-seq dataset. P-values calculated using two-tailed student’s t-test. E. Absolute values of log_2_ fold transcriptional changes in siDouble cells, of upregulated (yellow) vs downregulated (blue) cluster 2 genes from ATAC-seq dataset. P-values calculated using one-tailed student’s t-test. F. GO term analysis of up (yellow) vs down regulated (blue) cluster 2 genes in siDouble cells. Negative log_10_ p value of enrichment is represented on the y-axis.

The top 5 enriched GO pathways in cluster 2 genes are part of cell cycle and mitosis pathways while the other two clusters are involved in general metabolic and DNA damage pathways (Figure 3B). Based on our RNA-seq dataset, genes in cluster 2 demonstrate greater changes in transcription compared to cluster 1 genes (Figure 3C and 3D). In addition, cluster 2 corresponds to a subset of cell cycle related genes, including CENP-A, that are transcriptionally downregulated (Figure 3C). We then sought to examine the transcriptional changes of cluster 2 genes in the siDouble knockdown condition.

We separated the cluster 2 genes into highly upregulated and highly downregulated genes within the siDouble dataset. The analysis revealed that there is a higher degree of transcriptional change in the downregulated cluster 2 genes in the siDouble cells compared to the upregulated gene set (Figure 3E). Furthermore, GO term analysis of the two subsets shows that the upregulated cluster 2 genes in the siDouble cells are members of nucleotide synthesis pathways however, the downregulated cluster 2 genes are mitotic and cell cycle pathway members (Figure 3F). These enriched pathways in this subset of genes match the GO terms enriched in the total siDouble transcriptomics data set (Figure 2E) suggesting that these FOXM1 targets are regulated at the chromatin level by CENP-F. Taken together, this suggests that CENP-F and FOXM1 coordinately regulate chromatin accessibility at G2/M genes to drive their transcription.

### CENP-F regulated FOXM1-MMB complex formation

Given the evidence that CENP-F has a regulatory function at G2/M genes, we sought to understand the role of CENP-F on chromatin recruitment of FOXM1 by the MuvB-MYBL2 complex. FOXM1, MYBL2 and CENP-F are similarly upregulated in cancer cells based on the TCGA Pan-Cancer database (Figure 4A)[43]. Consistent with FOXM1 recruitment by MuvB, FLAG immunoprecipitation in FLAG-FOXM1 RPE cells pulls down MYBL2 and LIN54, a member of the MuvB complex. To determine how CENP-F contributes to MuvB complex function using siRNA-mediated knockdown and observed that loss of CENP-F weakens the association of MYBL2 and LIN54 with FOXM1 (Figure 4B and 4C). Taken together this supports the hypothesis that CENP-F is a regulator of the formation of FOXM1-MuvB complex which sits at promoters of CEN/KT genes.

**Figure 4.**
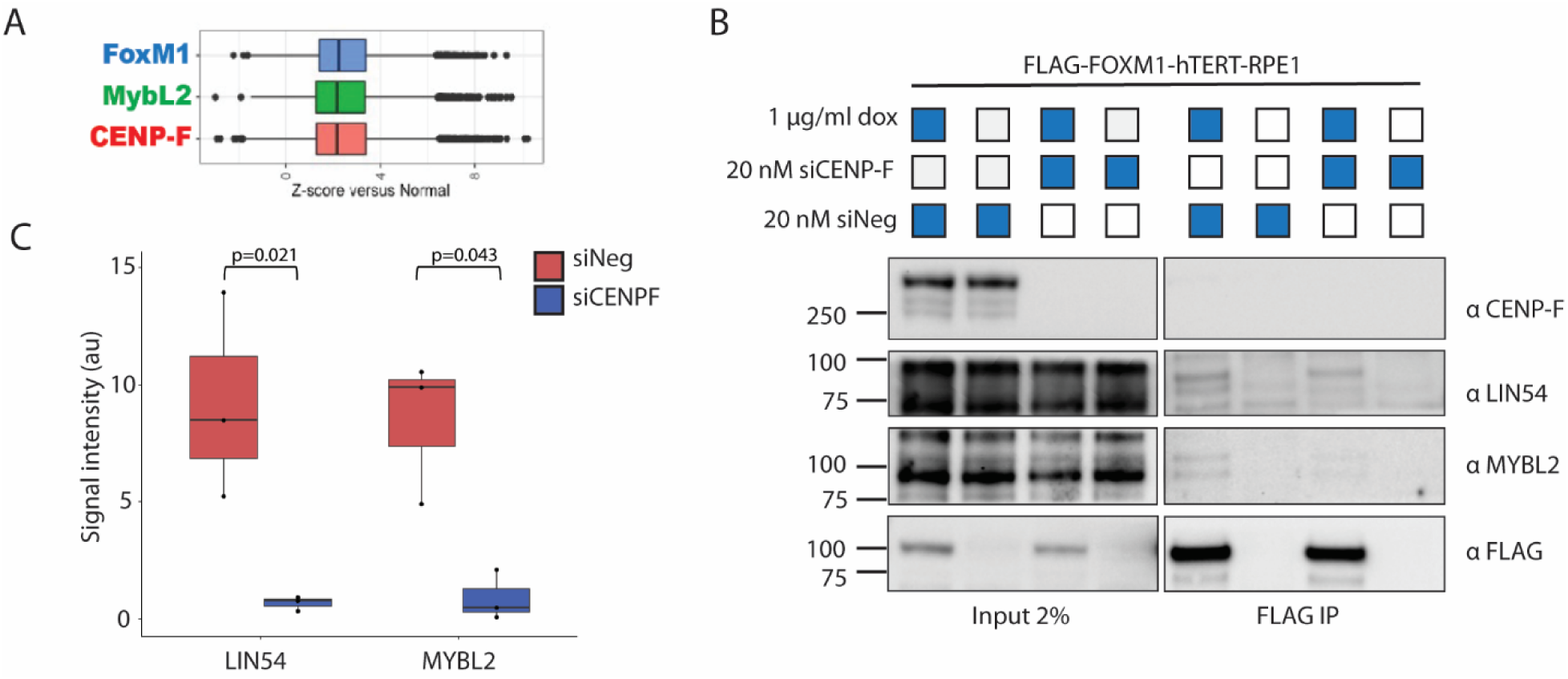
CENP-F regulates FOXM1-MMB complex formation. A. TCGA pan cancer analysis of gene expression represented as z-score of cancer vs normal of FOXM1, MYBL2 and CENP-F. B. Immunoblot of FLAG immunoprecipitation in FLAG-FOXM1-hTERT-RPE1 cells. Cells were treated with doxycycline for 48 hours to induce FOXM1 expression and simultaneously treated with 20nM siNeg or siCENP-F. C. Quantification of immunoprecipitation of LIN54 or MYBL2 shown in (B). P-values calculated using two-tailed student’s t-test. N=3.

### FOXM1 and MYBL2 drive activation of a specific subset of CEN/KT proteins

Previous literature has demonstrated coordinated misregulation of centromere and kinetochore (CEN/KT) proteins in several cancer types[24, 25]. To examine this systemic misregulation in cancer cells compared to normal cells, we analyzed expression data from the TCGA Pan-Cancer Atlas. Consistent with previous data, we observed that the majority of centromere and kinetochore (CEN/KT) genes are upregulated in cancers, across of cancer types and origins (Figure 5A). Notably, there are distinct exceptions to the upregulation namely CENP-C and MIS12 which demonstrate limited change in expression in cancer samples compared to normal tissues.

**Figure 5.**
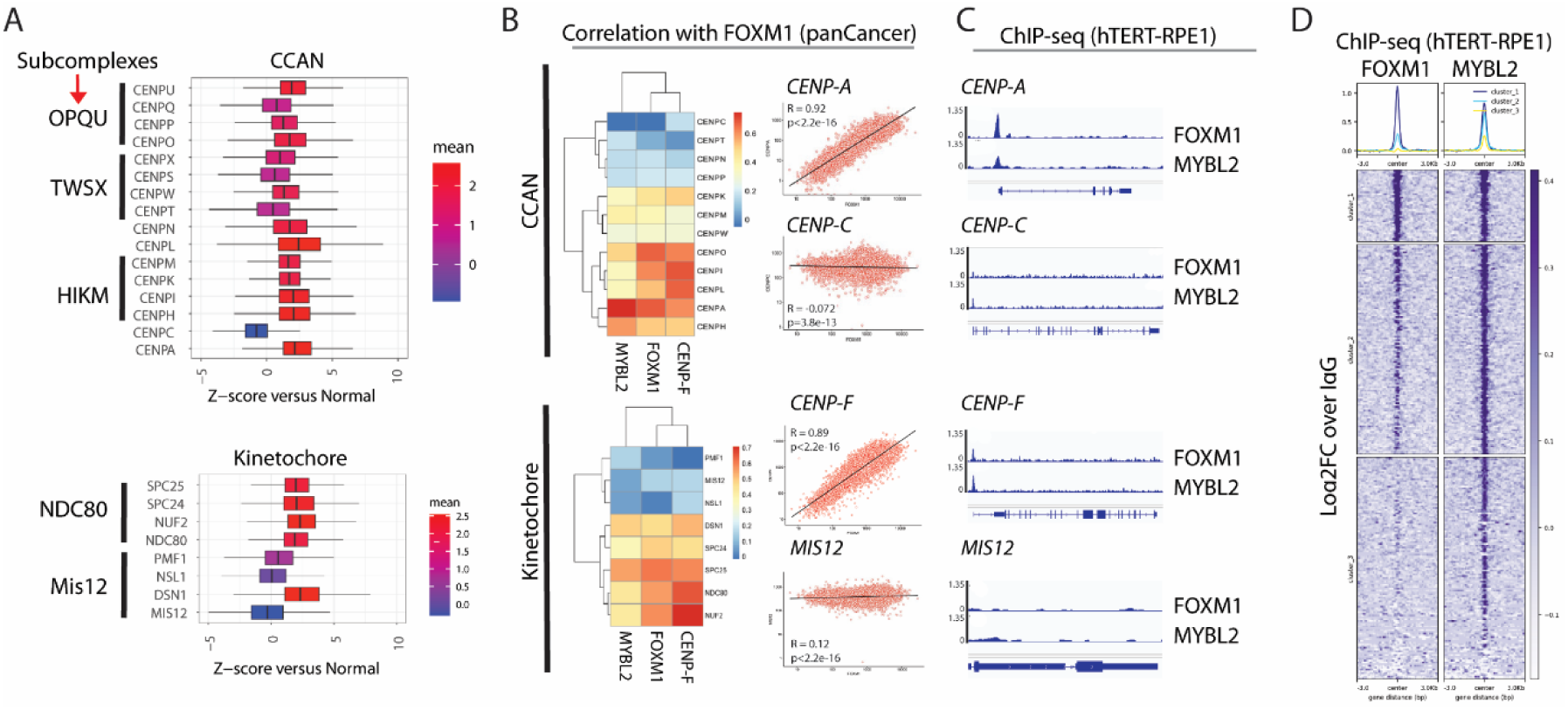
FOXM1 and MYBL2 co-occupy a specific subset of CEN/KT proteins. A. TCGA pan cancer analysis of gene expression represented as z-score of cancer vs normal of CEN/KT subcomplexes. B. Correlations of CEN/KT components with FOXM1 from TCGA data. Scatterplot of select genes shows degree of correlations between FOXM1 and CENPA or CENPC of the CCAN complex and FOXM1 and CENPF or MIS12 of the kinetochore complex. Pearson’s correlation coefficient was calculated from z-scores of cancer versus normal. C. FOXM1 and MYBL2 ChIP profiles in hTERT-RPE1 cells showing peaks at selected genes. D. Metaplot of FOXM1 and MYBL2 log_2_FC ChIP signal vs IgG at FOXM1-MYBL2 intersected consensus peaks with k-means clustering = 3.

To identify drivers of CEN/KT transcription, we examined correlations of mitotic transcription factors FOXM1 and MYBL2 with the genes using the TCGA Pan-Cancer database (Figure 5B). Consistent with the previously established role of these transcription factors, they show strong correlation with the CEN/KT genes. Specifically, while CENP-A and CENP-F show strong correlation with FOXM1 (and MYBL2, data not shown), the genes that fail to show upregulation in cancer such as CENP-C and MIS12, are not correlated. Next, we examined FOXM1 and MYBL2 occupancy at promoters of the CEN/KT genes using ChIP-seq in non-transformed hTERT-RPE1 cells. Peaks called in our ChIP-seq reveal that FOXM1 and MYBL2 strongly co-occupy promoters, correlated genes such as CENP-A and CENP-F show strong peaks, but occupancy at the uncorrelated ones such as CENP-C and Mis12 is limited (Figure 5C and 5D).

### FOXM1 and CENP-F driven regulation is differential in cancer cells

We observed that in normal cells, FOXM1 overexpression or knockdown did not result in a significant change in CENP-F levels. Thus, we hypothesized that that the FOXM1-CENP-F regulatory loop may differ in the cancer cells compared to normal cells. To that end, we treated a breast cancer cell line (MCF7) and a prostate cancer cell line (PC-3) with siFOXM1 for 48 hours and examined the protein levels of CENP-F in comparison to the hTERT-RPE1 cells. Loss of FOXM1 leads to an insignificant increase of CENP-F protein expression in the normal hTERT-RPE1 cells, compared to the expected loss of CENP-F in both the cancer lines (Figure 6A and Figure 6B). Changes in CENP-F protein levels are consistent with CENP-F transcript changes in the normal compared to cancer cells in response to FOXM1 loss (Figure 6C) suggesting that the differential expression is certainly on the RNA level. This is consistent with the repression phenotype seen in the FOXM1 overexpression hTERT-RPE1 cell (Supplemental Figure 2E) and suggests that FOXM1-CENP-F mediated transcriptional overactivation may be specific to the context of cancer.

**Figure 6.**
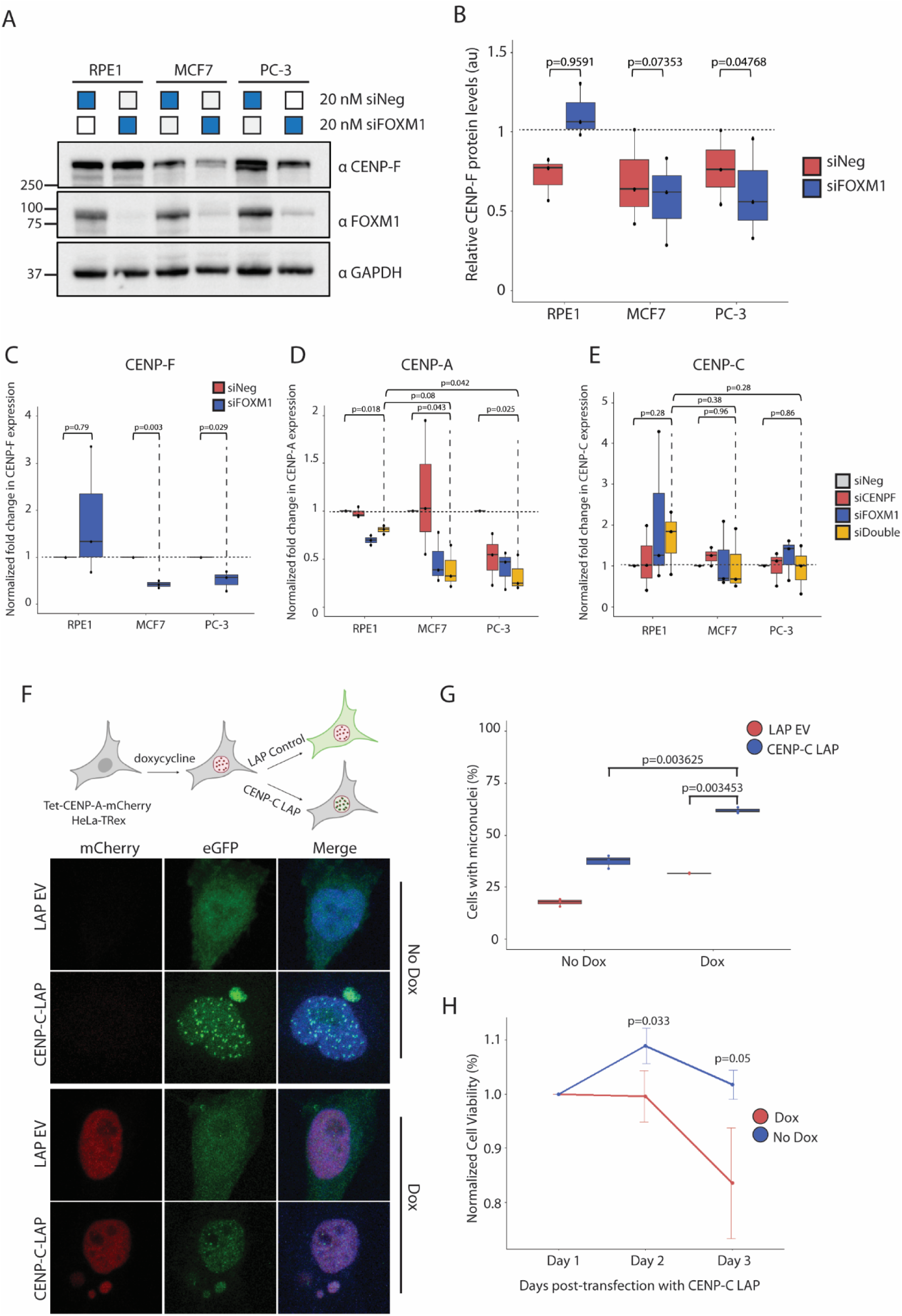
FOXM1-CENP-F cancer-specific regulation excludes CENP-C to promote survival in cancer cells. A. Immunoblotting of hTERT-RPE1, MCF7 and PC-3 cells treated with 20nM siNeg or siFOXM1 for 48 hours. B. Quantification of CENP-F protein levels in response to siNeg or siFOXM1 in normal vs cancer cells lines shown in (A), normalized to GAPDH. P-values calculated using one-tailed student’s t-test. N=3 C. RT-qPCR of CENP-F transcript levels, normalized to GAPDH, in FOXM1 knockdown conditions across normal and cancer cell lines. P-values calculated using two-tailed student’s t-test. N=3. D. RT-qPCR of CENP-A transcript levels, normalized to GAPDH, in knockdown across normal and cancer cell lines. P-values calculated using two-tailed student’s t-test. N=3. E. RT-qPCR of CENP-C transcript levels, normalized to GAPDH, in knockdown conditions across normal and cancer cell lines. P-values calculated using two-tailed student’s t-test. N=3. F. Representative images of Tet-CENP-A-mCherry HeLa-Trex cells induced for 48 hours and transfected with either CENP-C LAP or LAP empty vector (EV). Imaging was conducted for mCherry (CENPA), eGFP (LAP) and DAPI. G. Quantification of percentage of cells with micronuclei in each condition shown in (E). P-values calculated using two-tailed paired student’s t-test. N=3. H. Percent cell viability calculated by growth curves over 72 hours post transfection in each condition. P-values calculated using two-tailed paired student’s t-test. N=3.

### Escape of CENP-C from FOXM1 regulation as a potential cancer cell survival mechanism

Based on evidence that FOXM1-MYBL2 may only regulate a subset of genes and that CENP-F-FOXM1 regulation may be cancer cell specific, we examined differential transcriptomic effects of coordinated loss of CENP-F and FOXM1 in the normal and cancer cell lines. While the double CENP-F and FOXM1 knockdown leads to a 26.5% loss of CENP-A RNA levels in hTERT-RPE1, the double knockdowns in the MCF7 and PC3 lead to a 66.93% and 66.97% loss of CENP-A RNA respectively. This is a significant loss compared to both the parental cancer cells and the hTERT RPE1 cells (Figure 6D). Interestingly, CENP-C levels are largely unaffected in any condition (Figure 6E).

The differential regulation of CENP-A and CENP-C by FOXM1 and MYBL2 may be attributed to a cancer cell survival mechanism that works to limit chromosome instability levels. To determine if CENP-C overexpression biases cells towards catastrophic chromosome misegregation and cell death when CENP-A is overexpressed, we expressed LAP (eGFP) tagged CENP-C in doxycycline inducible CENPA-mCherry HeLa-TRex cells (Figure 6F). We found that double overexpressing cells showed a significantly higher percentage of micronuclei compared to either CENP-A or CENP-C only overexpressing cells (Figure 6G). Furthermore, cells overexpressing both CENP-A and CENP-C experienced reduced viability after 72 hours compared to either protein upregulated alone, suggesting that cells with elevated levels of CENP-A and CENP-C result in higher cell death (Figure 6H). Taken together, this suggests that FOXM1 and MYBL2, regulated by CENP-F, drive a specific subset of the CEN/KT genes to promote increased proliferation in cancer cells while preventing catastrophic chromosome segregation.

## Discussion

Driven by errors in chromosome segregation, chromosome instability (CIN) provides an opportunity for cancer cells to evolve tumorigenic abilities. Mitotic transcription factors FOXM1 and MYBL2 are proposed to drive the expression of genes involved in the transition from G2 to M, including the components of the centromere and kinetochore (CEN/KT). Coordinated misexpression of the CEN/KT is prevalent in cancer and is correlated with FOXM1 expression. While FOXM1 is proposed to be the key driver of mitotic transcription, the mechanism of context specific changes in this transcriptional program in normal versus cancer cells is poorly understood. Previous literature has proposed a synergistic relationship between FOXM1 and CENP-F, an outer kinetochore protein, in cancer cells, however there is currently limited mechanistic explanation for this observation [38]. Previous reports have suggested the involvement of CENP-F in transcription such as the association with ATF4 and the potential regulatory function with RB1. Our study expands the understanding of CENP-F function in transcription by providing evidence of the chromatin altering function of CENP-F at G2/M genes[40, 41].

We hypothesized that cancer-specific changes in the FOXM1 program, through CENP-F regulation of chromatin accessibility, lead to misregulation of mitotic genes and subsequent segregation errors. We observe that our established subset of CEN/KT genes, for example CENP-A but not CENP-C, is differentially coordinately regulated by CENP-F and FOXM1 specifically in the context of cancer but not normal cells. We determined that FOXM1 occupies the promoters of a subset of CEN/KT genes with MYBL2. Determining the role of CENP-F in FOXM1-driven transcription and how this function changes in the context of cancer is important for developing strategies to target chromosome instability phenotypes in cancer cells.

Previous experiments presume that the centromere and kinetochore genes are uniformly regulated by FOXM1, however, our analysis shows that a subset of centromere and kinetochore genes are uncoupled from FOXM1 regulation. This is apparent in both the analysis of TCGA expression data, and the lack of an effect of FOXM1 knockdown on CENP-C expression. CENP-C acts as a key lynchpin between the CENP-A nucleosome and the CCAN complex[44, 45]. CENP-A overexpression results in the deposition of CENP-A nucleosomes into non-centromeric loci via the chaperone DAXX[21, 46]. Without coordinated upregulation of CENP-C, the CENP-A nucleosomes in the chromosome arms may be limited in their ability for ectopic centromeres, leading to chromosome misegregation. Indeed, we show that overexpression of CENP-A and CENP-C together leads to significantly higher micronuclei formation and reduced cell viability. Therefore, the selective regulation of the CCAN without CENP-C may buffer cancer cells from a fatal level of chromosome instability.

Co-silencing of FOXM1 and CENP-F using siRNA-mediate knockdown has a coordinated and systemic effect on chromosome segregation. We determined a cluster of FOXM1 bound G2/M genes, including CENP-A, that experience changes in chromatin landscape in response to loss of CENP-F. These genes are coordinately downregulated in double silenced cells. Considering there is no direct association between CENP-F and FOXM1 proteins, CENP-F is likely co-regulating transcription with FOXM1 at the chromatin level. Interestingly, our work demonstrated complex formation between FOXM1, MYBL2 and LIN54 and this interaction is weakened by loss of CENP-F.

Previous studies suggest that the MYBL2-MuvB complex recruits FOXM1 to the CHR motifs at promoters of G2/M genes[12]. After MYBL2 is evicted, the FOXM1-MuvB complex drives transcription at these sites and loss of CENP-F in prostate cancer cells has shown to reduce FOXM1 occupancy at these promoters[38]. Our data in non-transformed hTERT-RPE1 cells treated with RO-3306 shows a specific but statistically insignificant trend – loss of CENP-F results in decreased FOXM1 occupancy and increased MYBL2 occupancy. This suggests that CENP-F may be involved in regulating proper occupancy of the FOXM1-MuvB complex at promoters but the drastic phenotype in response to the loss of CENP-F may be a cancer-context specific effect.

Based on evidence that CENP-F is differentially regulated in cancer cells and the context dependent changes in the FOXM1-MYBL2 transcriptional program, we hypothesize that perhaps, in mitotic cancer cells, CENP-F promotes FOXM1 occupancy and then stimulates MYBL2 eviction at promoters at G2/M transition, ultimately allowing for activation of mitotic genes. A previous study demonstrates that CDK1 is required for FOXM1-MMB activation, however it is also required during S-phase for late origin firing during replication. Thus, the cell balances CDK1 levels via ATR-mediated CHK1 activation, to prevent excess expression leading to premature mitotic entry [11, 47]. Though it promotes G2 arrest, RO3306 is a CDK1 inhibitor and thus, we are likely unable to confirm our hypothesis due to insufficient activation and incomplete occupancy of FOXM1 and MYBL2.

CENP-F is a unique component of the kinetochore because it is involved in the transcriptional regulation of its own complex. This begs the question if this is a feedback loop and if CENP-F can transmit information from the kinetochore to influence expression of the CEN/KT components. As FOXM1 cycles through activator and repressive phases during the cell cycle, we propose a similar model for CENP-F in that it contributes to balancing mitotic gene expression with FOXM1. Perhaps, when unbound, CENP-F associates with FOXM1 binding sites in order to increase transcription and when sufficient CENP-A is produced and the centromere can assemble, CENP-F returns to the kinetochore to perform its canonical function. Additionally, our observation that the double knockdown has increased chromosome instability phenotype compared to the individual knockdowns and we can interpret this as a coordinated transcriptomic effect on G2/M genes. However, we cannot ignore that the CENP-F affect could be due to the direct loss at the kinetochore. An important next step in our study would be to produce CENP-F binding mutants to allow parsing of the canonical versus non-canonical functions of the protein, especially in the context of cancer. Developing DNA binding mutants of CENP-F would contribute to delineating the two potentially distinct contributions of the CENP-F protein by allowing us to prevent the transcriptional function of CENP-F at G2/M promoters and then study how this may affect FOXM1-mediated transcriptional activation.

Overall, we show that FOXM1 regulation of G2/M is systemic, and CENP-F coordinately contributes by regulating chromatin accessibility and promoting FOXM1-MuvB complex formation. We provide a mechanistic explanation for the coordination of CENP-F and FOXM1 transcription and evidence for cancer-specific changes in FOXM1-MMB activation of CEN/KT genes, regulated by CENP-F. Not only do we illuminate a novel function of CENP-F, we provide insight into the complexity of CEN/KT misexpression and subsequent chromosome misegregation.

## Materials and Methods

### Cell culture and stable cell line production

hTERT-RPE1 PAC KO cells were cultured in DMEM/F12 with 10% Tet-tested fetal bovine serum (FBS) and 1% penicillin-streptomycin (P/S). MCF7 and HEK293T cells were cultured in DMEM with 10% FBS and 1% P/S. PC-3 cells were cultured in RPMI-1640 media with 10% FBS and 1% P/S. Cells were incubated at 37°C with 5% CO_2_ and passaged with 0.05% trypsin.

Tet-FLAG-FOXM1 hTERT-RPE1 stable cells line were produced using lentiviral transduction. Lentivirus was produced by transfecting 2×10^6^ HEK293T cells on a 10cm plate using 12ug of target plasmid encoding FLAG-FOXM1c, 6ug of psPAX2 viral packaging vector and 3ug pMD2.G viral envelope vector overnight with Lipofectamine 3000 (Invitrogen) according to manufacturer protocol. Culture medium was collected 48 hours and 72 hours post-transfection and passed through at 0.45µM SCFA filter. hTERT-RPE1 PAC KO cells were transduced with virus for 24 hours at an MOI of 1 with 8ug/mL of polybrene, followed by 3 ug/mL puromycin selection. Tet-mCherry-CENP-A HeLa-TRex cells were produced as previously described[48]. Tet-inducible cells lines were doxycycline induced for 48 hours at 2ug/mL in conjunction with any other treatments.

### siRNA and LAP plasmid transfection

For siRNAs, cells were transfected with 20nM ON-TARGETplus pools (Dharmacon) using RNAiMAX Transfection Reagent (Invitrogen) according to manufacturer protocol. 10nM of each siRNA was used for the double knockdown conditions. Cells were collected 48 hours post-transfection.

The LAP empty vector plasmid was a gift from Todd Stukenberg and the CENP-C-LAP plasmid was a gift from Iain Cheeseman produced as previously described [49]. 2.5×10^5^ Tet-mCherry-CENP-A HeLa-TRex cells were transfected with 1ug of either CENP-C-LAP or empty vector plasmid in 6-well plates with Lipofectamine 3000 (Invitrogen) according to manufacturer protocol. Cells were collected 48 hours post-transfection and doxycycline induction.

### Immunofluorescence

Cells were treated and then plated in culture medium on coverslips pre-coated with 0.1% poly L-lysine. The coverslips were washed in PBS, fixed in 4% paraformaldehyde for 10 minutes and quenched with 100mM Tris-HCl pH8. Fixed cells were then incubated in a blocking buffer of PBS with 0.1% Triton X-100, 2% FBS, 1.6mg/mL BSA for 1 hour at room temperature. Primary antibodies were diluted in blocking buffer as follows: CENPF (Mouse 1:400), FOXM1 (Rabbit 1:100), a-Tubulin (Mouse 1:1000), ACA (Human 1:50). After washing, cells were incubated with fluorophore conjugated secondary antibodies (Cy3 and FITC) at 1:1000 for 1 hour. Lastly, cells were incubated with 0.2ug/mL DAPI in PBS for 5 minutes and mounted on glass slides with Prolong Gold. Z-stacks were acquired using a 63X oil-immersion objective lens on a Zeiss LSM800 microscope. Collected images were maximum projected and channels were scaled identically within panels.

Nuclear signal analysis was conducted using a custom ImageJ2/Fiji script as previously described[50]. To calculate percentage of cells with micronuclei, total cells per condition were counted along with cells showing evidence of micronuclei based on DAPI channel. Lagging anaphase events were calculated using a publicly available ImageJ2/Fiji script using standard parameters (https://github.com/Elowesab/elowelab/blob/master/Lagging_chromosomes_2020-10-06.ijm)[51].

### Immunoblotting

Whole-cell lysates were prepared in radioimmunoprecipitation assay (RIPA) buffer, and total protein concentration was quantified using a bicinchoninic acid (BCA) assay kit. Lysates were denatured in 1X Laemmli buffer with 6.5% βME and boiled at 95°C for 15 minutes. Equal mass of cell lysate was loaded in a 4-12% gradient SDS-PAGE precast gel and run at 100V for 1.25 hours. Gels were transferred to nitrocellulose membranes using wet transfer in 20% methanol at 350mA for 1.5 hours. Membranes were then blocked in 5% BSA and then analyzed using desired primary and secondary antibodies. Western blots were developed using a BioRad ChemiDoc imager and then quantified by densitometry using ImageJ2/Fiji gel analysis feature.

### Immunoprecipitation

For FLAG immunoprecipitation in FLAG-FOXM1 hTERT-RPE1 cells, cells were doxycycline induced for 48 hours. Cells were lysed in NETN buffer (20mM Tris-HCl pH 7.5, 150mM NaCl, 0.5% NP-40, 0.5mM EDTA, and 1X Protease Inhibitors) for 30 mins at 4°C with gentle agitation and then incubated with 20U benzonase and 10mM MgCl_2_ to break down chromatin for 30 mins at 37°C. The lysate was clarified for 15 minutes at 4°C at maximum speed. 5% input was removed and stored, and the remainder of the lysate was incubated with pre-washed anti-FLAG M2 magnetic beads (Sigma) overnight at 4°C. The lysate-conjugated FLAG beads were then washed 5x in NETN buffer and eluted with 7.5ug 3X FLAG Peptide in NETN buffer. Lysates were the denatured in 1X Laemmli buffer with 6.5% βME and purified proteins were boiled at 95°C for 15 minutes. 1% input and the full volume of the IP in each condition, were immunoblotted with anti-LIN54, anti-MYBL2, anti-FLAG and anti-GAPDH antibodies as described above. Quantification of band intensities was conducted using the gel analysis tool in ImageJ2/Fiji.

### Real-time quantitative PCR

Total RNA was extracted from pre-treated cells using the RNeasy Mini Kit (Qiagen) according to the manufacturer protocol. Extracted RNA was quantified using a nano-spectrophotometer and then stored at -80°C. 500ng of RNA was converted to cDNA using iScript™ cDNA Synthesis Kit (Bio-Rad) according to the manufacturer protocol. The cDNA is then further diluted 1:5 and qPCR reactions were set up as follows: 2.5µl diluted cDNA, 5µl of iTaq Universal SYBR Green Supermix, 0.5µl of 10µM reverse primer, 0.5µl of 10µM forward primer and 1.5µl of ddH2O per well. qPCR samples were run in technical duplicates in a 96 well plate and detected using Bio-Rad CFX Connect Real-Time PCR System using the following program: 95°C for 30 seconds and then 40 cycles of 95°C for 5 seconds and 60°C for 30 seconds.

### Cell Proliferation Assay

1.5×10^4^ Tet-mCherry-CENPA cells were plated per well in a 24 well format and transfected 24 hours later with 100ng of LAP empty vector or CENP-C LAP per well. After 24 hours of transfection, cells were counted using a ViCell Blu and the viability percentage of each well was recorded. Cells were then induced with doxycycline for 48 hours and viability percentage of each well was recorded every 24 hours. The viability of CENP-C LAP transfected cells was normalized against LAP empty vector transfection wells. Cells were plated in technical duplicates and experiment was repeated for three biological replicates.

### RNA-seq

Total RNA was extracted from cells treated as outlined above. The NEBNext® Poly(A) mRNA Magnetic Isolation Module (New England Biolabs) was used as per manufacturer protocol to isolate mRNA for input into the NEBNext® Ultra™ II RNA Library Prep Kit (New England Biolabs) using NEBNext Unique Dual Index Primers (New England Biolabs). Library preparation was conducted as per manufacturer protocol. Libraries were quality checked using Agilent D1000 TapeStation kit and Qubit Flourometer. FLAG-FOXM1-hTERT RPE1 experimental samples were sequenced on an Illumina NextSeq (Shilatifard lab) to generate 2×75 bp paired-end reads and FOXM1 and CENP-F siRNA experimental samples were sequenced on an Illumina HiSeq (Admera Health) to generate 2×150 bp paired-end reads. We obtained 40M reads per sample. Data analysis including read demultiplexing, alignment and differential expression were done using the Ceto pipeline (https://github.com/ebartom/NGSbartom) with standard parameters. Gene Set Enrichment Analysis (GSEA) of differentially expressed genes was performed using the Molecular Signature DataBase from the Broad Institute[52, 53]. TCGA Pancancer Atlas datasets were downloaded from cBioPortal (https://www.cbioportal.org) as mRNA expression z-scores relative to normal samples (log RNA Seq V2 RSEM). Downstream analyses were conducted using R.

### ChIP-seq and ChIP-qPCR

ChIP was performed in hTERT-RPE1 cells as follows: 50×10^6^ cells were crosslinked with 1% paraformaldehyde and 1% FBS in PBS for 10 min at room temperature, followed by quenching with 0.2M glycine. Cells were then washed in PBS, and flash frozen in liquid nitrogen. Cells were then lysed and sonicated with the Covaris E220 with the following parameters: 600 seconds, 10% duty cycle, 140 peak intensity and 200 cycles per burst. The lysates were centrifuged and input across the samples was normalized by DNA concentration. 1% of the Input was removed and then DNA-protein crosslinked complexes were immunoprecipitated overnight at 4°C with the following antibodies: FOXM1 (4ug CST), MYBL2 (4ug Proteintech) and Rabbit IgG (4ug CST) along. Complexes were then bound to Protein A Dynabeads (Invitrogen) for 4 hours at 4°C and washed in lysis buffer. Reverse crosslinking reaction using proteinase K was performed at 65°C overnight, followed by DNA purification using Qiagen PCR purification kit. DNA libraries were produced using NEBNext Ultra II DNA Library Preparation kit and NEBNext Unique Dual Index Primers (New England Biolabs). Libraries were quality checked using Agilent D1000 TapeStation kit and Qubit Flourometer and then sequenced on the Illumina NovaSeq X (Admera Health) with 2×150 bp paired-end reads. We obtained 50M reads per sample. Data analysis including alignment of reads and peak calling was conducted using the nf-core/chipseq pipeline (https://github.com/nf-core/chipseq) [54]. Downstream analysis was performed using bedtools, deeptools and R. For ChIP-qPCR, ChIP was performed as described above with the following modifications: 12.5×10^6^ cells and 2ug of all antibodies were used. After DNA purification, ChIP DNA was subjected to RT-qPCR as outlined above. RO-3306 treatments were conducted in siRNA-treated cells for 16 hours at 9µM before ChIP-qPCR.

### ATAC-seq

ATAC-seq was conducted as described[55, 56] on siRNA-treated hTERT-RPE1 cells. Briefly, 5×10^4^ cells were isolated in cold 1xPBS and lysed with 0.1% NP-40, 0.1% Tween-20 and 0.01% Digitonin. Cells were then centrifuged to separate the nuclear material and the cytoplasm. The nuclear pellet was resuspended in transposition reaction mix using the Tagment DNA Enzyme and Buffer Kit (Illumina) as per manufacturer protocol. Transposition was conducted at 37°C for 30 minutes and DNA was then purified using Qiagen MinElute Reaction Cleanup Kit.

For library preparation, purified transposed DNA was PCR amplified using Nextera (Illumina) i5 common adapter and unique i7 index adapters and NEBNext High Fidelity 2x PCR Master Mix for 5 cycles with the following program: 72°C for 5 minutes, 98°C for 30 seconds, 98°C for 10 seconds, 63°C for 30 seconds and 72°C for 1 minute. The partially-amplified libraries were then analyzed by qPCR to determine additional PCR cycles with the goal of stopping amplification prior to saturation and avoid variation among samples due to PCR bias. Using 10% of the partially-amplified library, Nextera primers matching those used in the first PCR step and SYBR Green I (Invitrogen), the following qPCR program was performed: 98°C for 30 seconds, 98°C for 10 seconds, 63°C for 30 seconds and 72°C for 1 minute. The number of additional cycles needed for each sample was determined by calculating the number of cycle needed to reach 1/3 of the maximum RFU in that sample. PCR on the remaining 90% of the libraries was then conducted for the appropriate number of cycles per sample. Double-sided bead purification using AMPure XP beads was conducted to remove primer dimers and large >1000 bp fragments. Libraries were quality checked using Agilent D1000 TapeStation kit and Qubit Flourometer then sequenced on the Illumina HiSeq (Admera Health) with 2×150 bp paired-end reads. We obtained 70M reads per sample. Data analysis including alignment of reads and peak calling was conducted using the nf-core/atcseq pipeline (https://github.com/nf-core/atacseq) [54]. Downstream analysis was performed using bedtools, deeptools and R. FOXM1 ChIP-seq peaks from MCF7 used for the metaplots were obtained from ENCODE (https://www.encodeproject.org/experiments/ENCSR000BUJ/).

## Key Reagents

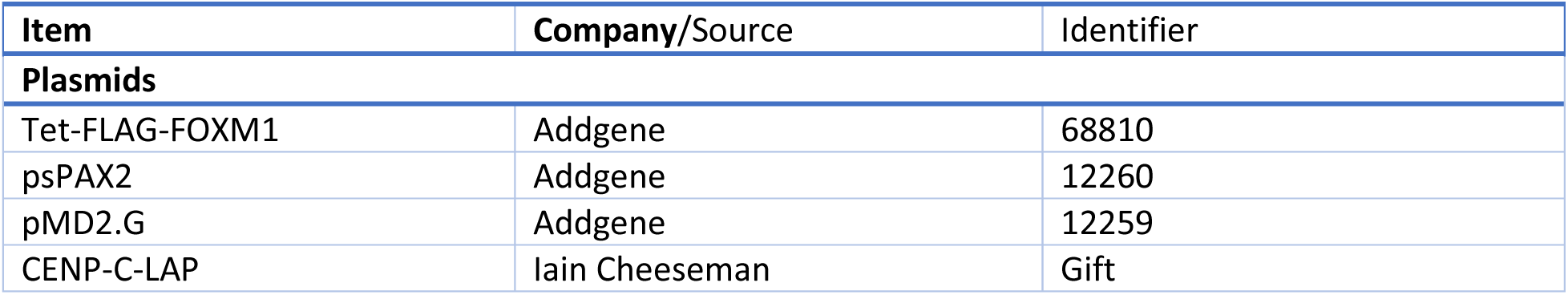
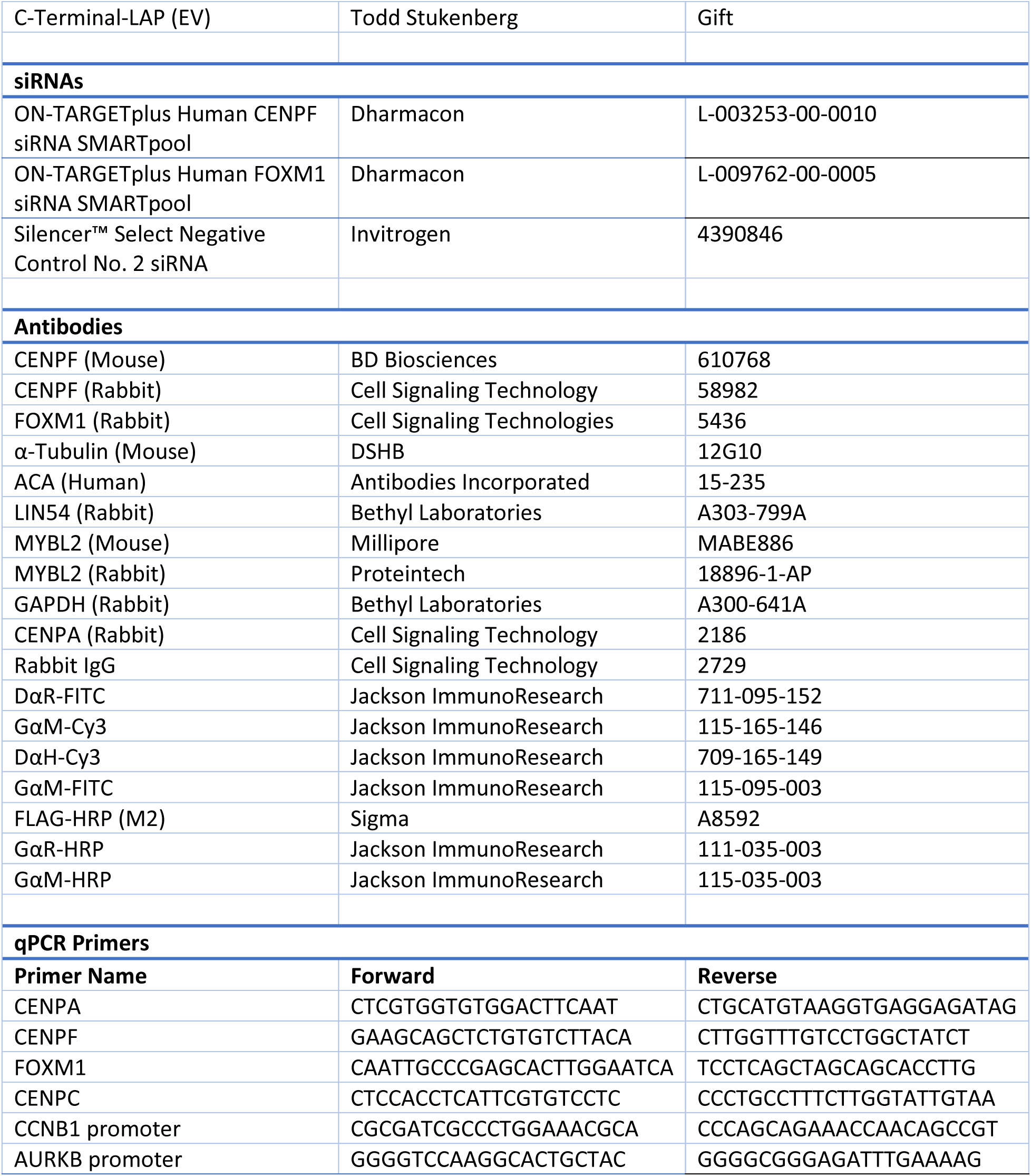

## Acknowledgements

We thank the Shilatifard Lab including Emily Rendleman, Marta Iwanaszko and Siddharta Das for their contributions to genomics experiments. We thank Elizabeth Bartom and Ann Hogan for assistance with genomic analysis. We thank the Foltz Lab members for their comments on the manuscript. D.R.F. was supported by NIH grants R01GM111907 and U01CA260699.

