## Supplemental Figures for "Contribution of CENP-F to FOXM1-mediated discordant centromere and kinetochore transcriptional regulation"

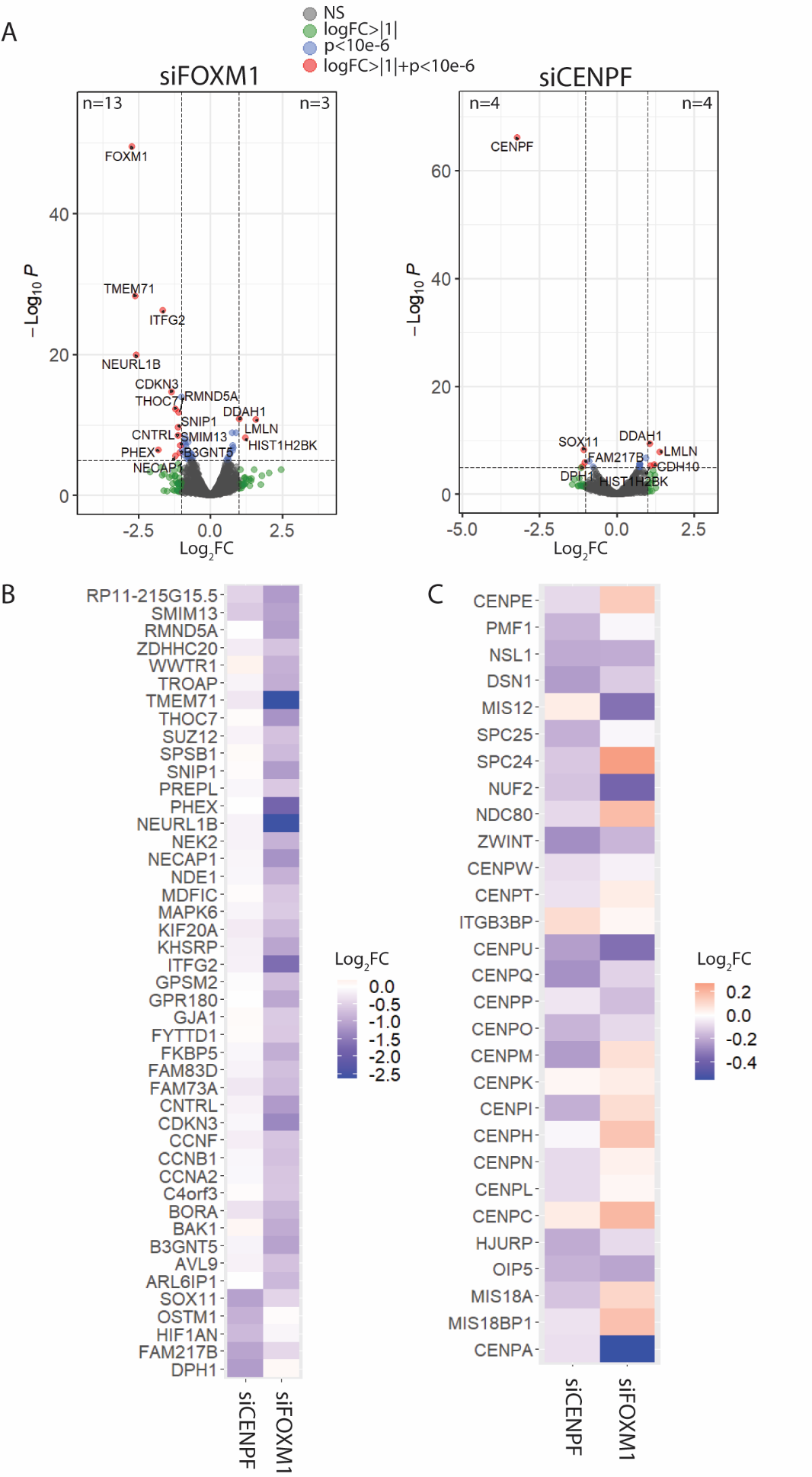

### **Supplementary Figures**

#### Supplementary Figure 1. FOXM1 is more transcriptionally active compared to CENP-F

A. Volcano plot showing distribution of genes from RNA-seq in siCENP-F and siFOXM1 treated hTERT-RPE1 cells. Genes in red have a log_2_FC>|1| and p-value>10e-6, genes in green have log_2_FC>|1|, genes in blue have p-value>10e-6 and grey dots represent non-significant genes (n=3).
B. Heatmap showing most significantly downregulated genes in both conditions.
C. Heatmap showing log_2_FC of CEN/KT genes in both conditions.

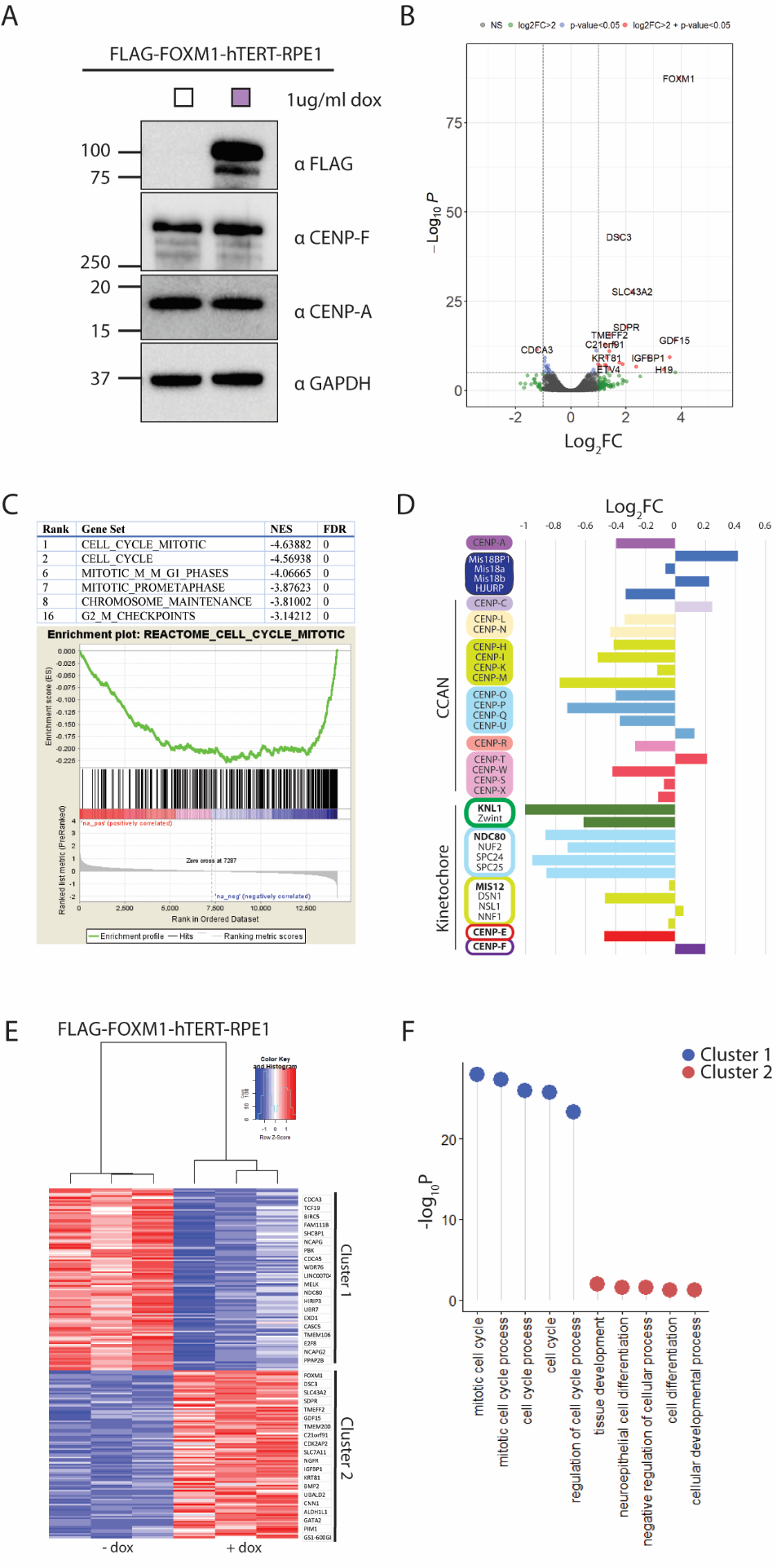

#### Supplementary Figure 2. FOXM1 is insufficient to drive transcription of CEN/KT genes and instead, represses G2/M genes and activates developmental pathways in asynchronous hTERT-RPE1.

A. Immunoblotting of FOXM1-FLAG-hTERT-RPE1 cell lines showing doxycycline induction.
B. Volcano plot showing distribution of genes from RNA-seq in FLAG-FOXM1-hTERT-RPE1 cells. Genes in red have a log_2_FC>|2| and p-value>0.05, genes in green have log_2_FC>|2|, genes in blue have p-value>0.05 and grey dots represent non-significant genes (N=3).
C. Gene set enrichment analysis using pre-ranked gene and log_2_FC list. Table shows list of mitotic pathways enriched in the top 20 gene sets. Example enrichment plot shows negative enrichment of top gene set ‘cell cycle mitotic.’
D. Bar graph showing log_2_FC of CEN/KT genes in FLAG-FOXM1-RPE RNA-seq dataset.
E. Heatmap showing clustering of most significantly downregulated (cluster 1) and upregulated (cluster 2) genes in FLAG-FOXM1-hTERT RPE1 RNA-seq dataset.
A. GO term analysis of cluster 1 and cluster 2. Negative log_10_ p value of enrichment is represented on the x-axis.

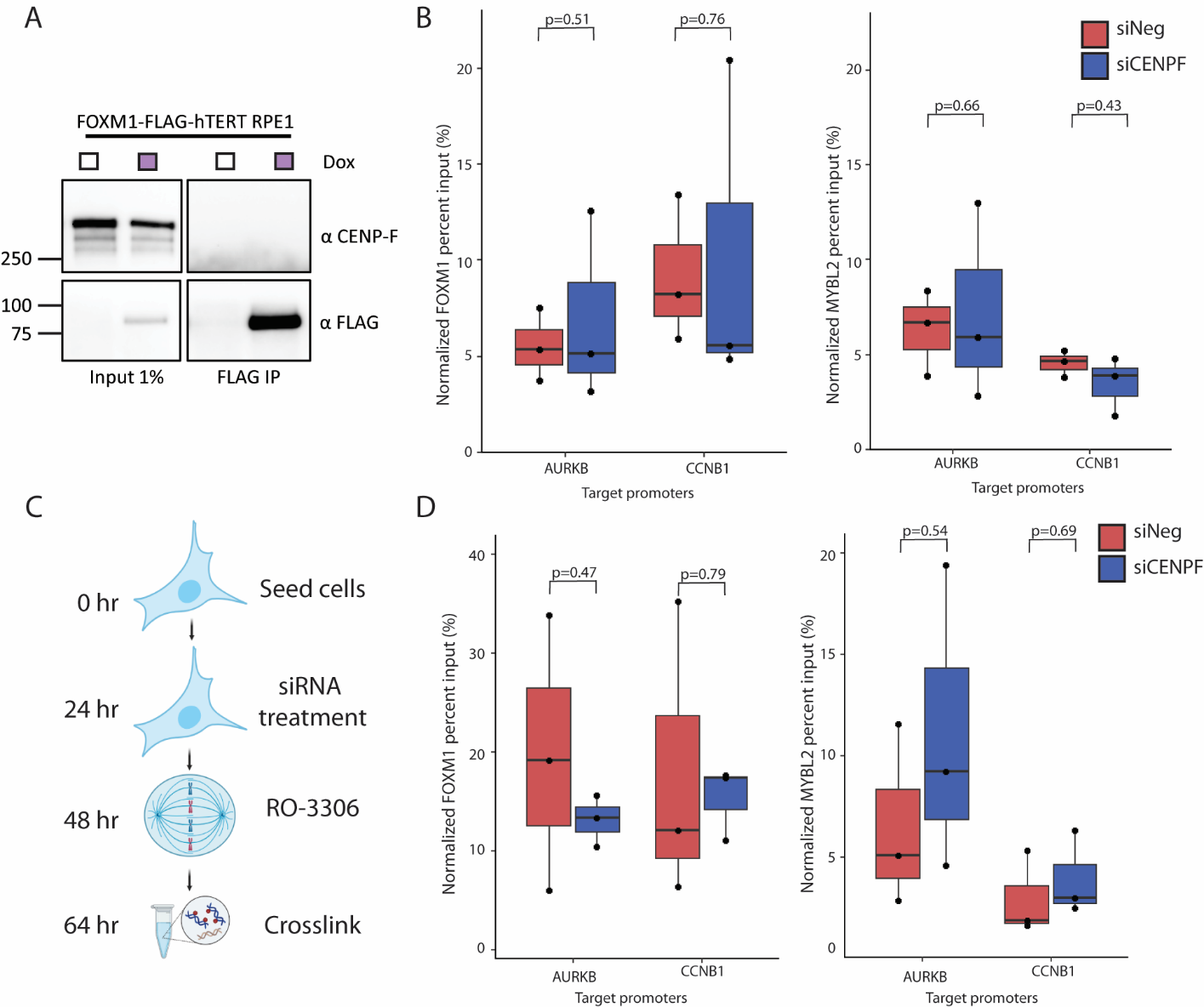

#### Supplementary Figure 3. Promoter occupancy of FOXM1 and MYBL2 at G2/M genes in CENP-F siRNA treated asynchronous cells.

A. Immunoblot of FLAG immunoprecipitation in FLAG-FOXM1-hTERT-RPE1 cells. Cells were treated with doxycycline for 48 hours to induce FOXM1 expression and input and IP were blotted back for CENP-F. N=3.
B. Percent input of FOXM1 (left) and MYBL2 (right) at AURKB and CCNB1 promoters in asynchronous hTERT-RPE1 treated with 20nM siNeg or siCENP-F for 48 hours. P-values calculated using two-tailed student’s t-test. N=3.
C. Schematic of RO-3306 treatment to enrich G2/M cells.
D. Percent input of FOXM1 (left) and MYBL2 (right) at AURKB and CCNB1 promoters in RO3306 treated hTERT-RPE1 cells treated with 20nM siNeg or siCENP-F for 48 hours. P-values calculated using two-tailed student’s t-test. N=3.
